# QUANTIFYING GLYCOGEN AND LIPID DROPLET SYNTHESIS IN OVARIAN AND CERVICAL CANCER CELLS USING DEUTERATED RAMAN PROBES WITH STIMULATED RAMAN SCATTERING MICROSCOPY

**DOI:** 10.64898/2026.03.16.712189

**Authors:** Ryan N. Pierson, Shovit A. Gupta, Mingyuan Zhang, Lucas C. kaiser, L. Nathan Tumey, Fake Lu

## Abstract

Epithelial ovarian cancer remains one of the most lethal malignancies among women, with late-stage diagnoses yielding 5-year survival rates below 30%. The metabolic heterogeneity of the tumor microenvironment (TME) highlights the need for methods capable of rapid, chemically specific phenotyping. Stimulated Raman scattering (SRS) microscopy when combined with deuterium labeled metabolites enables the non-invasive high contrast interrogation of cellular metabolic pathways. In this study, we used SRS microscopy to profile fatty acid and glycogen metabolism in epithelial ovarian cancer (SKOV-3) and cervical cancer (HeLa) cell models. Deuterium labeled glucose revealed striking differences in glycogen synthesis and intracellular distribution, with SKOV-3 cells exhibiting markedly greater single-cell heterogeneity than HeLa. Complementary measurements of lipid droplet (LD) synthesis and turnover under nutrient starvation further revealed cell-line-specific metabolic strategies, identifying LD and glycogen dynamics as a potential diagnostic marker of cancer metabolic phenotypes. These results demonstrate that SRS microscopy in the Raman silent region, paired with metabolic labeling, can sensitively resolve metabolic diversity across cancer cell subpopulations. Such metabolic phenotyping may inform both early diagnostic strategies and therapeutic approaches that combine cytotoxic treatment with targeted metabolic disruption.

## 1. Introduction

Dysregulated lipid and glucose metabolism along with frequent metabolic reprogramming are known hallmarks of cancer progression. Cancer cells will preferentially utilize aerobic glycolysis to support the formation of biomass and forego the process of oxidative phosphorylation (OXPHOS), a phenomenon known as the “Warburg effect” [1]. The metabolic plasticity seen in cancer imparts a challenging task in the development of effective anti-tumor drugs, with many patients suffering from a refractory period during treatment due to the diversity in cancer cell phenotypes [2, 3]. This plasticity can be observed at the single cell level and dynamically adjusts in response to changes in the tumor microenvironment (TME) [4]. TME has elevated levels of lactic acid, poor tissue oxygenation, and low glucose availability owing to aggressive glycolytic activity and cancer associated vasculature [5]. Despite these harsh conditions, proliferation of cancerous cells can continue partially due to the improved adaptability of metabolic reprogramming confers. Increased expression of fatty acid translocases, fatty acid-binding proteins, and free fatty acid receptors is frequently observed in cancer and collectively facilitates increased exogenous fatty acid (FA) uptake [6]. De novo FA synthesis can also occur in parallel with exogenous fatty acid uptake through conversion of citrate to palmitic acid [7]. The resulting palmitic acid is subsequently able to interact with elongases and stearoyl-CoA desaturase (SCD) to produce monounsaturated fatty acids (MUFAs). Acyl-CoA synthetase (ACS) then converts the intracellular MUFAs to acyl-CoA, and after a process of mitochondrial translocation, acyl-CoA is converted to acetyl-CoA via beta fatty acid oxidation (β-FAO) for immediate use in the TCA cycle. Alternatively, MUFA derived acyl-CoA can be converted into neutral lipids, such as triacylglycerols (TAGs), through Lipin and DGAT1/2 mediated reactions and packaged into lipid droplets [8].

The cellular intake of exogenous glucose in cancer is mediated by GLUT family transporters frequently overexpressed in cells undergoing Warburg metabolism [9]. Glucose carbon bond fate once inside the cell is complex and typically involves multiple metabolic pathways regulated within a single cell [10]. After 6C phosphorylation catalysis by hexokinases, glucose can be converted to pyruvate via aerobic glycolysis or polymerized in the form of glycogen via glycogenesis. Pyruvate as a metabolic intermediate acts at the crossroads between entrance into the TCA cycle, or conversion into lactate mediated by lactate dehydrogenase [11]. Should pyruvate be used with regards to the former possibility, it can contribute to *de novo* lipid droplet synthesis through TCA-cycle derived citrate conversion into palmitic acid.

Lipid droplets are a cellular organelle once only thought of as simple energy reservoirs for cells to tap into, and have gained much attention in recent years due to their now known multifunctionality in cancer cell survival [12]. Lipid droplet accumulation has been implicated as a characteristic of aggressive forms of certain cancer types, as they confer cytoprotective properties to a cell in the presence of anti-cancer drugs and other sources of cellular stress [13]. Current research on lipid metabolism in cancer often utilizes high specificity chemical profiling techniques such as mass spectrometry or high-performance liquid chromatography, which results in the destruction of the sample [14, 15]. Raman scattering is an inelastic scattering effect between incident light and the chemical bonds between atoms, which consequently enables the acquisition of chemical information from a sample.

Stimulated Raman scattering (SRS) microscopy can be used to acquire a chemical fingerprint of a sample in addition to the spatial location of chemical bonds in any given field of view. SRS microscopy differs from other Raman techniques, such as spontaneous Raman confocal spectroscopy or coherent anti-Stokes Raman spectroscopy (CARS) and offers some unique advantages over them. For instance, SRS does not suffer from the non-resonant background observed in CARS or the long acquisition times needed for resolving spontaneous Raman confocal microscopy [16]. Additionally, spectral data obtained through SRS is straightforward to analyze, as the intensity measured is directly proportional to the number of chemical bonds. The chemical bond contrast of biological samples when using SRS can further be enhanced by utilizing the “silent” region of the Raman spectrum located between 1900 cm^-1^ and 2700 cm^−1^. Chemical bonds with resonance in the silent region are not endogenous to biological tissues, minimizing cellular background when using compounds with chemical bonds that reside within this special domain [17]. Previous work has demonstrated the feasibility of using alkyne tags, which have strong resonance within the silent region, to visualize lysosome lipid droplet sequestration [18]. In this paper, deuterated oleic acid and glucose are used to spatially track and quantify the aberrant metabolism of each metabolite in ovarian and cervical cancer cells using SRS microscopy.

## 2. Materials and Methods

### 2.1. Cell culture and treatment

Both SKOV-3 and HeLa cell lines were acquired from the ATCC. For SKOV-3 cell culture, cells were grown in T25 flasks with McCoy’s 5a base medium supplemented with 10%/v fetal bovine serum (FBS) and 1%/v antibiotic-antimycotic (AB-AM) until approximately 80% confluence was observed. HeLa cells were grown in DMEM instead of McCoy’s 5a. Upon reaching confluence, cells were either transferred to 50 mm glass bottom dishes for glucose experiments, or 50 mm polystyrene dishes with inserted glass cover slips for oleic acid experiments. Oleic acid-d_34_ was supplemented to the respective complete cell culture medium at a final concentration of 100 μM for each cell line upon cell transfer to the polystyrene dishes. For glucose experiments, cells were supplemented with 25 mM D-glucose-d_7_ in glucose free DMEM with 10%/v FBS and 1%/v AB-AM. Cells were treated over 24h for oleic acid treatment and 72h for glucose treatment. For 24h and 48h oleic acid-d_34_ uptake comparisons, cells treated over 48h were denied medium renewal to observe differences in lipid droplet handling under nutrient scarce conditions. Verification of glycogen stores in cell samples was done through increasing concentrations of GS-1 inhibitor (MZ-101, MedChemExpress) with SKOV-3 cells given 25 mM D-glucose-d_7_ over 72h. For lipid droplet depletion experiments, the same concentration of oleic acid was used with increasing glucose deprivation time (0h, 24h, 48h) in culture to assess lipid droplet depletion dynamics. Oleic acid-d_34_ treated cells were fixed with 4% paraformaldehyde (PFA) in phosphate buffered saline (PBS) for 10 minutes and immediately imaged using a lab-built integrated SRS/two-photon fluorescence (TPF) microscope. D-glucose-d_7_ treated cells were imaged whilst alive to better retain labile glycogen within the cell bodies. For cell doubling time experiments, SKOV-3 cells were seeded at a cell density of 100,000 cells/ml in 6-well plates. Cells were given 25 mM D-glucose or D-glucose-d_7_ over a period of 72 hours with medium renewal every 24 hours. The final cell count was used with the initial cell count to determine the cell doubling time under each metabolic condition. All cell counting experiments were done using an automated cell counter (EVE, NanoEntek).

### 2.2. Spontaneous Raman confocal microscopy

Spectral acquisitions of SKOV-3 cells treated with 50 mM D-glucose or D-glucose-d_7_ for 72 hours were acquired using a spontaneous Raman confocal microscope (**Fig. 2(E-G)**, Renishaw InVia). Measurement parameters were set to the following: 20x objective and a 785 nm laser source at 100% power using the 1200 lines/mm diffraction grating. The cells were then centrifuged at 200*g for 5 minutes and the supernatant was discarded. To eliminate any traces of phenol red, cells were resuspended in PBS and centrifuged again at 200*g for 5 minutes. After discarding the supernatant once more, the cell pellets were then placed on a glass slide wrapped with polished aluminum foil to eliminate background signal.

**Fig. 1.**
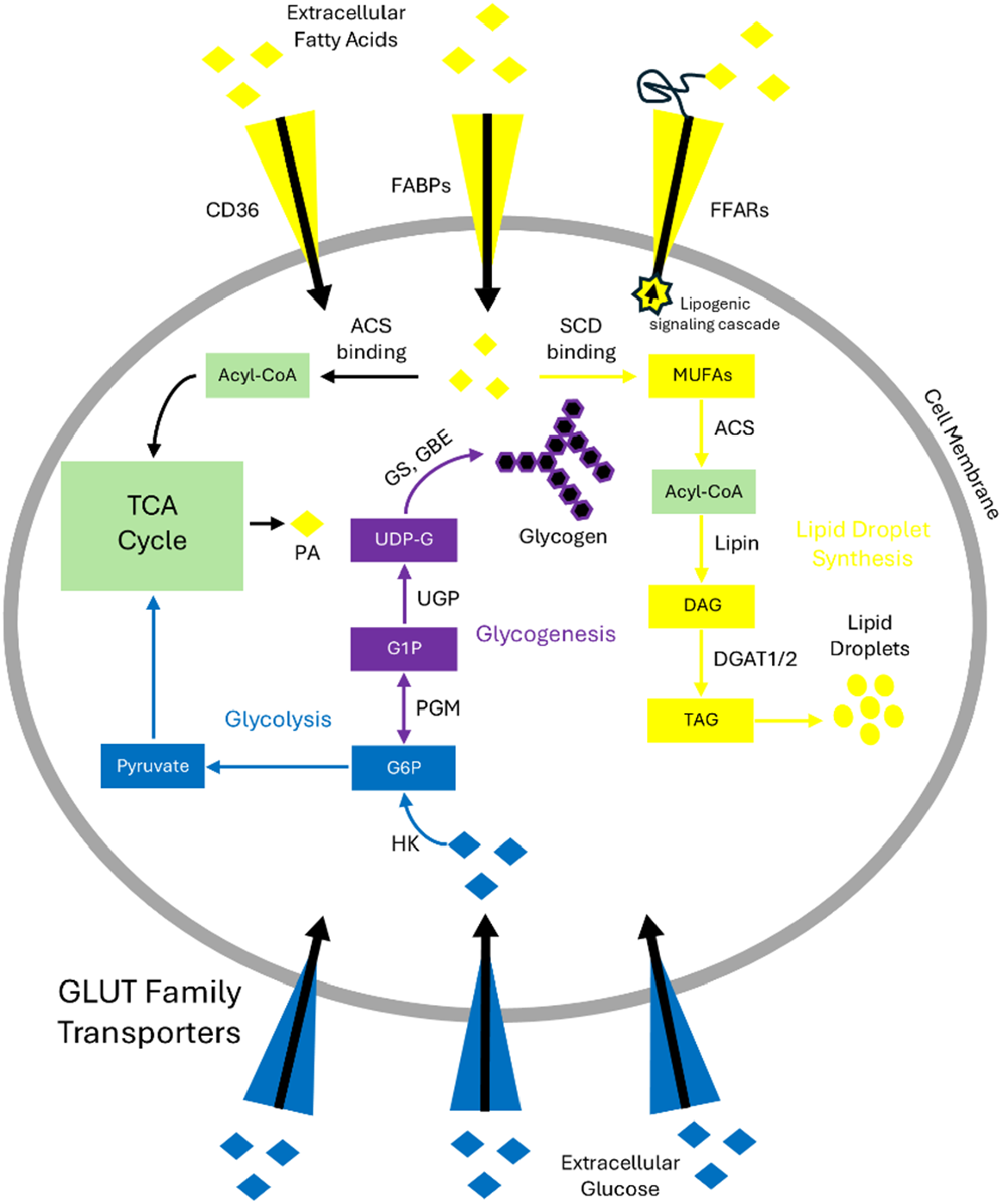
Simplified map of lipid and glucose metabolism in cancer. Extracellular fatty acids are transported to the cytosol via fatty acid transporters, such as CD36 and FABPs, while FFARs can upregulate lipogenic pathways through sensing of fatty acids via signaling cascades. Cytosolic fatty acids can be used as fuel for the TCA cycle after consecutive binding with related enzymes. Alternatively, fatty acids can bind with SCD, producing MUFAs. MUFAs can be further processed into TAGs and stored as lipid droplets after Lipin and DGAT1/2 mediated enzymatic reactions. Exogenous glucose is transported into the cell via glucose transporters, such as GLUT-1. Glucose is then phosphorylated by HK to produce G6P, which can be used in glycolysis, or act as a substrate for glycogenic enzymes during the process of glycogenesis. Abbreviations: CD36; platelet glycoprotein 4, FABP; fatty acid binding protein, FFAR; free fatty acid receptor, ACS; acyl-CoA synthetase, SCD; stearoyl-CoA desaturase, DGAT1/2; diacylglycerol acyltransferase-1/2, HK; hexokinase, PGM; phosphoglucomutase, UGP; UDP-glucose pyrophosphorylase, GS; glycogen synthase, GBE; glycogen branching enzyme, LDH; lactate dehydrogenase, PA; palmitic acid, MUFAs; monounsaturated fatty acids, DAG; diacylglycerol, TAG; triacylglycerol.

**Fig. 2.**
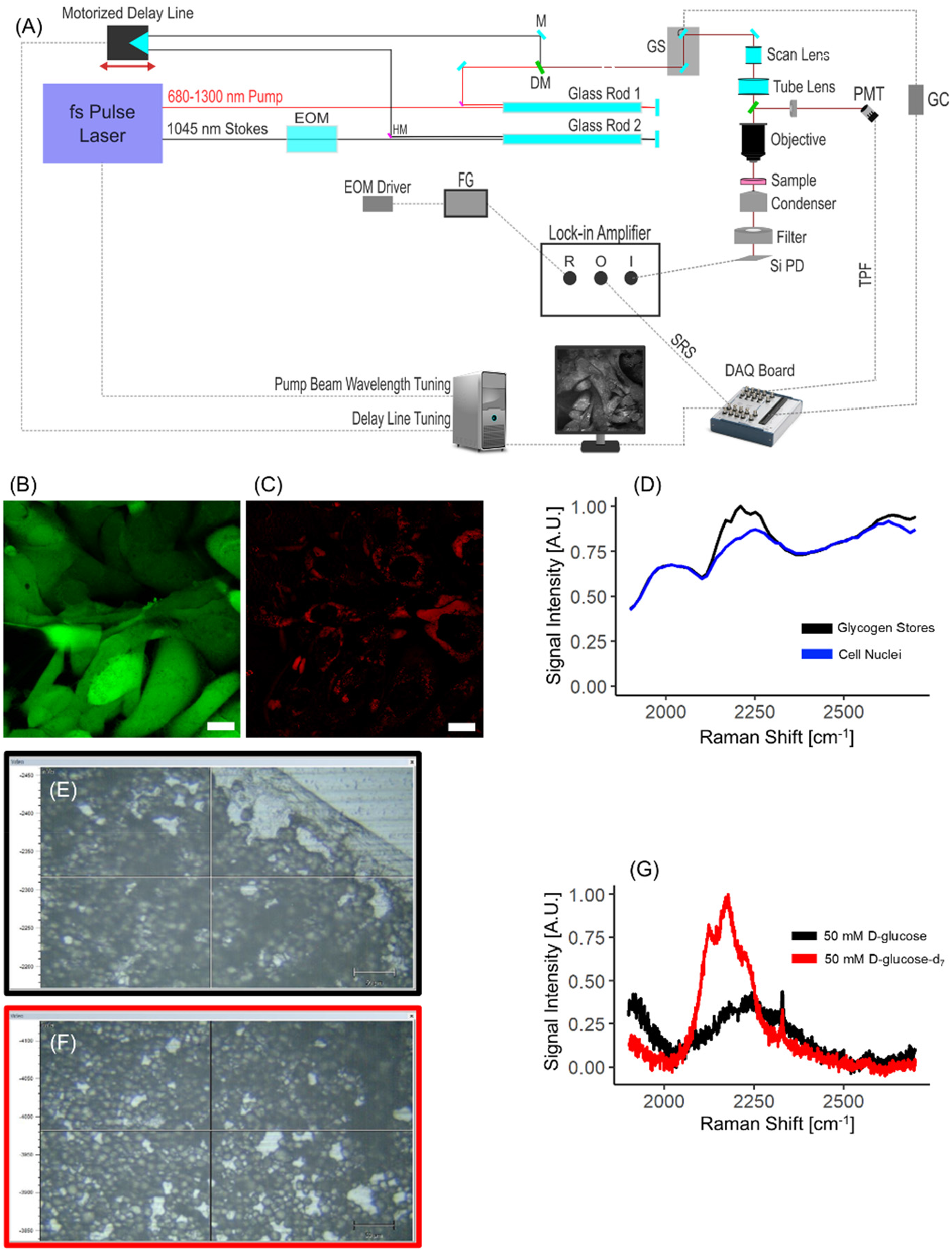
(A) Schematic view of the SRS imaging system integrated with two-photon fluorescence (TPF) microscopy. EOM: electro-optic modulator, PBS: polarizing beam splitter, M: mirror, DM: dichroic mirror, GC: galvo controller, GS: galvo scanner, FG: function generator, R: lock-in reference, O: lock-in output, I: lock-in input, Si PD: silicon photodetector. (B) Two-photon fluorescence of SKOV-3 cells stained with calcein-AM. (C) SRS image of SKOV-3 cells treated with 25 mM D-glucose-d7 over 72 hours at the carbon-deuterium resonance mode. Scale bar = 20 µm. (D) Hyperspectral SRS (1900-2700 cm^-1^) of fixed SKOV-3 cells treated with 25 mM D-glucose-d7 over 72h. (E) A representative field of view used for Raman confocal analysis of a SKOV-3 cell pellet treated with 50 mM D-glucose over 72h. (F) A representative field of view used for spontaneous Raman confocal analysis of a SKOV-3 cell pellet treated with 50 mM D-glucose-d_7_ over 72h. (G) Corresponding spectra covering the silent region for each cell pellet sample.

### 2.3. Hyperspectral SRS

At resonance and off-resonance Raman shifts for D-glucose-d_7_ were determined using hyperspectral SRS on fixed SKOV-3 cells that were treated with 25 mM D-glucose-d_7_ over 3 days. It was determined that the optimal carbon-deuterium (CD) peak and off-peak for D-glucose-d_7_ treated cells was 2209 cm^-1^ and 2321 cm^-1^, respectively. Raman shifts for the oleic acid-d_34_ peak and off-peak were previously determined to be 2112 cm^-1^ and 1991 cm^-1^, respectively.

### 2.4. Multi-channel SRS/TPF microscopy

Raman shifts measured within the high wavenumber region corresponded with CH_3_ (2930 cm^-1^) and CH_2_ (2852 cm^-1^) resonance modes. The CH off-resonance was measured at 2791 cm^-1^. Raman shifts of 2209 cm^-1^ and 2321 cm^-1^ were used for measuring the CD peak and off-peak of D-glucose-d_7_ treated cells, respectively. Raman shifts of 2112 cm^-1^ and 1991 cm^-1^ were used for measuring the CD peak and off-peak of oleic acid-d_34_ treated cells, respectively. The two-photon fluorescence component of the imaging system was used to determine cell viability in the presence of deuterated nutrients. Cell viability staining of live D-glucose-d_7_ treated SKOV-3 was performed using a 2 μM working solution of Calcein-AM in Hank’s Balanced Salt Solution (HBSS) followed by a 30-minute incubation period at 37°C and 5% CO_2_. The cells were then immediately imaged using the GFP emission filter of the TPF component of the integrated SRS/TPF system (optical path and imaging applications used are shown in **Fig. 2(A-D)**.

### 2.5. Statistical analysis and image processing

For CH_3_ and CH_2_ image background removal, the corresponding CH off-resonance image was subtracted from either resonance mode before image analysis. Similarly, CD off-peak measurements for both oleic acid-d_34_ and D-glucose-d_7_ treated cells were subtracted from the corresponding on-peak CD images. Image preprocessing of D-glucose-d_7_ images was done to correct the illumination gradient present at lower signal intensities. A Gaussian filter with a rolling ball radius of 45 was used on a duplicate CD image and subtracted from the original CD image before performing CD bond segmentation. Measurement of CD bond area fraction for oleic acid-d_34_ and D-glucose-d_7_ treated cells was done by dividing the total pixel count of the CD pixels by the total pixel count of the cell bodies. Cell body pixel count was determined through Huang thresholding of CH_3_ images. CD bond pixel count was determined using an automatic Python workflow. In brief, a grayscale image is further smoothed with a median blur to reduce high-frequency noise while preserving cellular boundaries. Following this denoising step, we applied Otsu’s global thresholding method to every image. If we assume that the image signal may be divided into two classes (background and signal), Otsu’s algorithm determines the threshold value that minimizes within-class variance, thereby separating background from signal without requiring manual adjustment. The Otsu-derived threshold was recorded for each image and used to generate two outputs: (i) a binary mask in which pixels above the threshold were classified as CD-positive, and (ii) a masked intensity image, in which CD-positive regions identified by (i) retained their intensity values, and all others were set to zero. For area fraction analysis, binary masks were used for pixel quantification. Statistical testing was done using either Student’s paired t-test or one-way ANOVA followed by a Tukey’s HSD post hoc test when applicable.

## 3. Results

### 3.1 SKOV-3 cell viability is maintained in the presence of deuterium analogues

In our pursuit to determine whether the incorporation of deuterium-labeled metabolites has any deleterious effect on cell health, we first assessed SKOV-3 ovarian carcinoma cell viability following treatment with D-glucose-d_7_. Live cell staining with calcein AM returned typical esterase activity and plasma membrane integrity (Fig. 3A). Robust fluorescence was observed throughout the cytoplasm of SKOV-3 cells exposed to D-glucose-d_7_, indicating that metabolic substitution with the deuterated sugar did not compromise overall cellular viability. Although strong calcein AM signal was broadly distributed, localized reductions in fluorescence intensity were observed in certain intracellular regions. These dimmer regions are consistent with acidic organelles, such as lysosomes, in which calcein fluorescence is known to be quenched due to pH sensitivity rather than loss of cell viability [19].

**Fig. 3.**
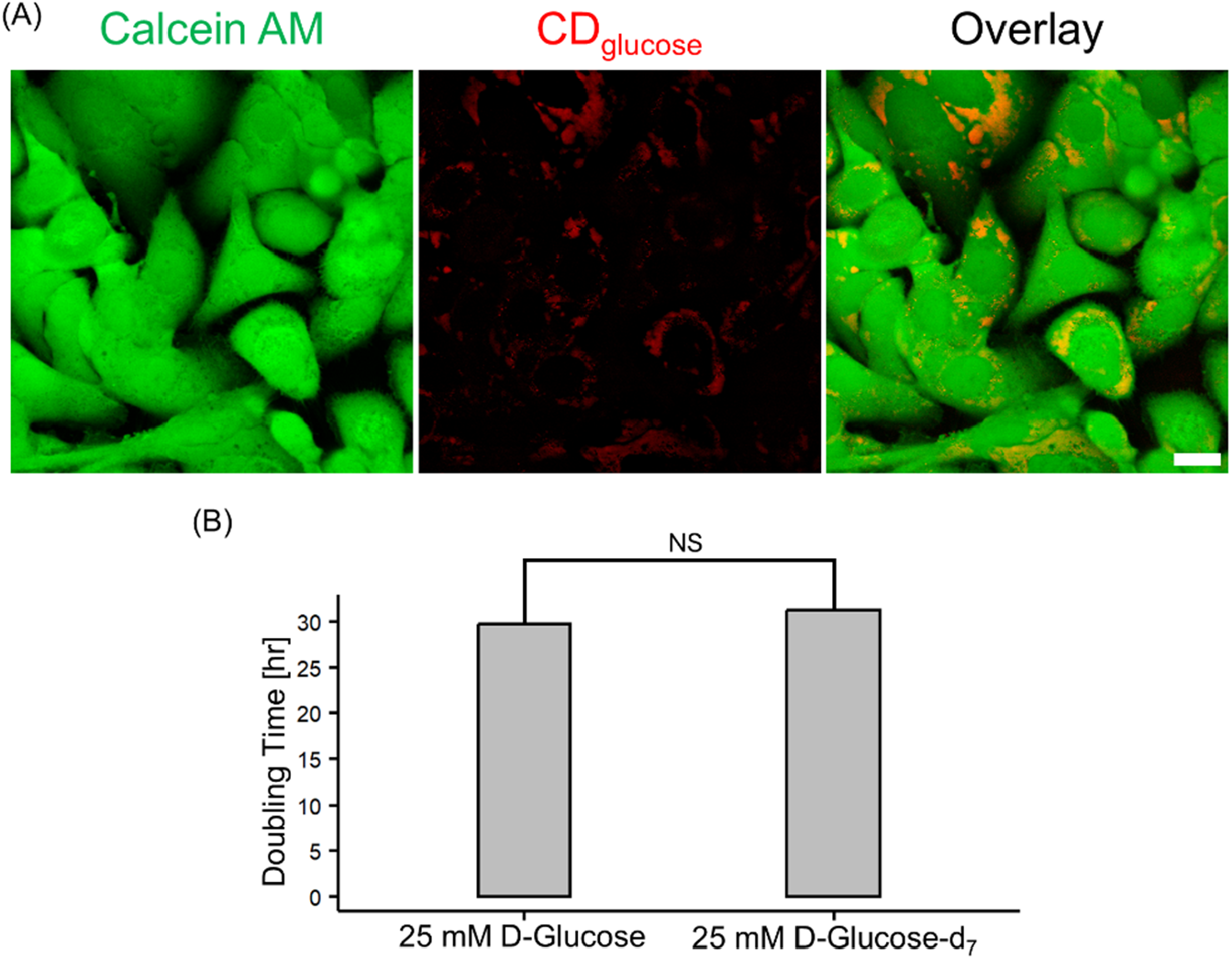
(A) TPF/SRS microscopy of live SKOV-3 cells treated with 25 mM D-glucose-d_7_ over 72 hours. Scale bar = 20 µm. (B) Cell doubling time measurement of SKOV-3 cells treated with either 25 mM D-glucose-d_7_ or D-glucose. N = 3 biological replicates. ‘NS’ denotes a non-significant result.

To further establish whether long-term exposure to deuterium analogues might alter cellular proliferation, we next quantified SKOV-3 doubling times comparing native D-glucose with D-glucose-d_7_ at a concentration of 25 mM. Cells cultured with the deuterated analogue displayed a modest increase in doubling time compared with the native glucose control group. However, this difference did not reach statistical significance (Fig. 3B). The absence of a measurable impairment in growth kinetics supports the interpretation that D-glucose-d_7_ substitution does not introduce overt metabolic stress to SKOV-3 cells.

The results from calcein AM viability staining and cell doubling time analysis demonstrate that the presence of D-glucose-d_7_ is well tolerated by SKOV-3 cells. These results reinforce the non-invasive nature of stable isotope labeling approaches for live-cell metabolite tracking and quantification, suggesting that deuterated glucose can be incorporated into cell culture systems without introducing confounding cytotoxic effects [20].

### 3.2 SRS imaging of deuterium-incorporated macromolecules in cells

To gain insight into how different carbon sources contribute to macromolecular storage in cancer cells, we supplemented cultures with deuterated analogues of either oleic acid or glucose. For lipid tracking and quantification, oleic acid-d_34_ treated cells were imaged at the CD stretching resonance (2112 cm^-1^) and its corresponding off-resonance mode (1991 cm^-1^), enabling specific visualization of deuterium-enriched lipid droplets (LDs) in live cells.

In both SKOV-3 ovarian carcinoma and HeLa cervical carcinoma cells, treatment with oleic acid-d_34_ resulted in strong CD signal localized to lipid droplets, with these droplets exhibiting a uniform spatial distribution throughout the cytoplasm (Fig. 4A). No discernible differences in droplet size or morphology were observed within a given field of view across either cell type, suggesting that exogenously supplied fatty acids were readily incorporated into neutral lipid stores in both cancer pathologies. These findings are consistent with prior reports that some cancer cells are highly permissive to extracellular lipid uptake and readily channel long-chain fatty acids into lipid droplet formation [21].

**Fig. 4.**
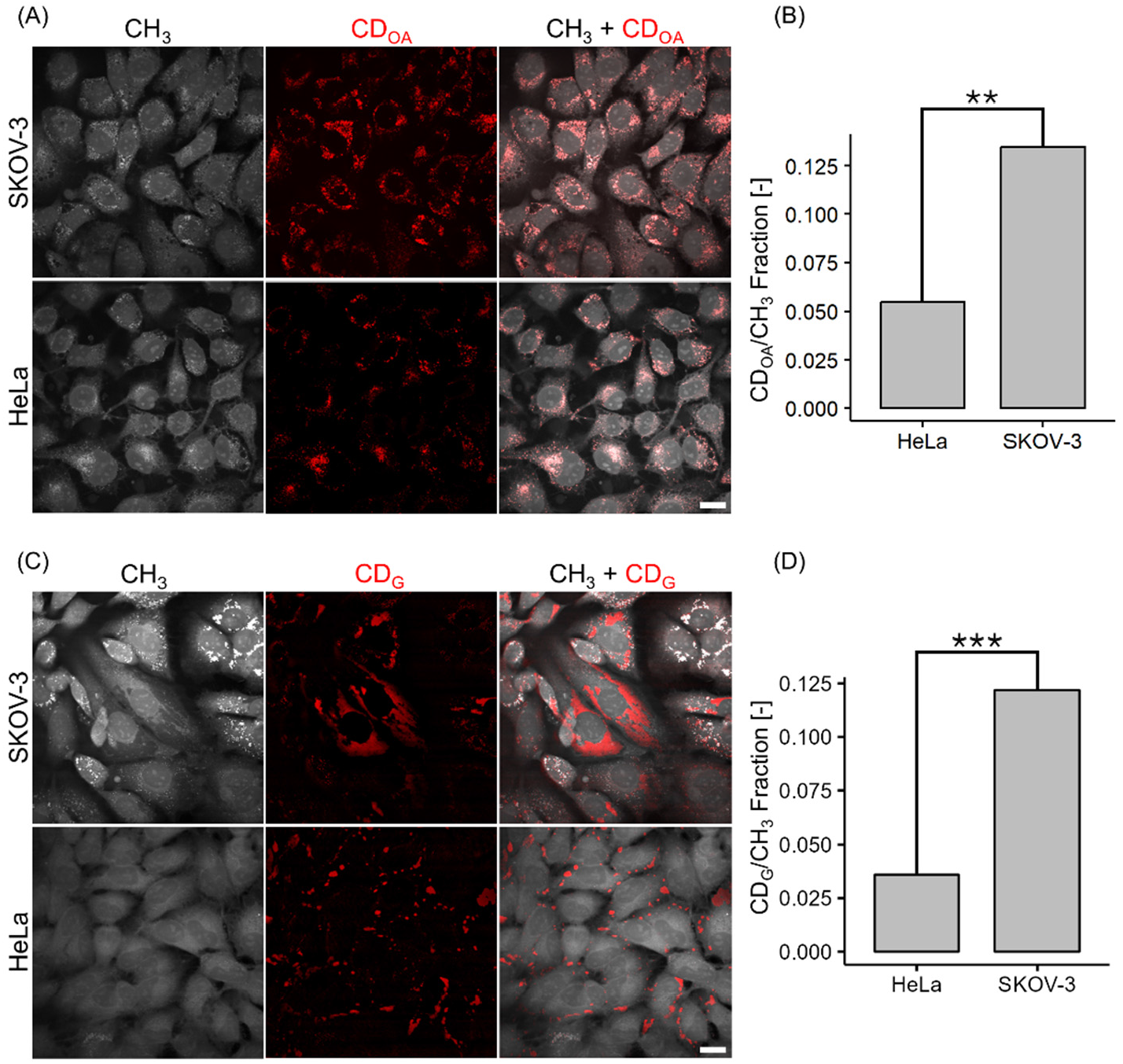
(A) SKOV-3 and HeLa cells treated with 100 µM oleic acid-d34 over 24 hours. (B) Comparison of oleic acid-d34 derived CD bond distribution between the two cell lines. (C) SKOV-3 and HeLa cells treated with 25 mM D-glucose-d7 over 72 hours. (D) Comparison of D-glucose-d7 derived CD bond distribution between the two cell lines. Scale bar = 20 µm. Statistical analyses were performed using Welch’s unpaired t-test. ‘^***^’ denotes a p value lower than 0.001. ‘^**^’ denotes a p value lower than 0.01. N = 3 biological replicates for oleic acid-d_34_ experiments. N ≥ 3 biological replicates for D-glucose-d_7_ experiments.

When quantifying lipid droplet accumulation, marked differences emerged between the two cell lines. SKOV-3 cells demonstrated a significantly greater capacity for lipid storage than HeLa cells under identical treatment conditions (Fig. 4B). This elevated lipid sequestration may reflect enhanced lipogenic or lipid storage programs in ovarian carcinoma cells, a phenotype previously linked to tumor aggressiveness and metabolic plasticity [22].

We next examined deuterated glucose utilization in the same cell lines. Unlike the homogeneous LD distributions observed with oleic acid supplementation, glucose-derived macromolecule synthesis exhibited strikingly different patterns. In SKOV-3 cells, glycogen accumulation following deuterated glucose treatment was heterogeneous, with individual cells showing variable levels of storage (Fig. 4C). In contrast, HeLa cells displayed a more uniform response, with glycogen appearing consistently across the population. Importantly, direct comparison of glycogen abundance revealed that SKOV-3 cells stored larger glycogen reserves relative to HeLa (Fig. 4D). These results suggest that while both cell types retain the capacity to divert glucose into glycogenesis, SKOV-3 cells exhibit greater metabolic heterogeneity, suggesting subpopulation-specific differences in nutrient handling and tumor cell adaptability.

Together, these findings highlight distinct metabolic strategies between SKOV-3 and HeLa cells. Both rely on exogenous fatty acids for lipid droplet formation, yet differ in efficiency of accumulation, and while both can direct glucose into glycogen, SKOV-3 cells display greater variability and overall abundance of glycogen storage. Such high variance in utilization of carbon sources across cancer pathologies underscores the importance of metabolic context in interpreting Raman-based isotope labeling studies of tumor models.

A further distinction between the two cancer models emerged when the dynamics of lipid droplet (LD) accumulation were monitored over extended culture periods. Following supplementation with 100 µM oleic acid-d_34_, SKOV-3 cells continued to accumulate deuterium-labeled lipid droplets even after 48 hours of incubation (Fig. 5). In contrast, HeLa cells initially formed LDs but exhibited a subsequent decline in droplet number under the same conditions. While found to be statistically insignificant, this divergent pattern between the two cell lines suggests that SKOV-3 cells favor long-term lipid retention, whereas HeLa cells engage in more active lipid droplet hydrolysis and recycling.

**Fig. 5.**
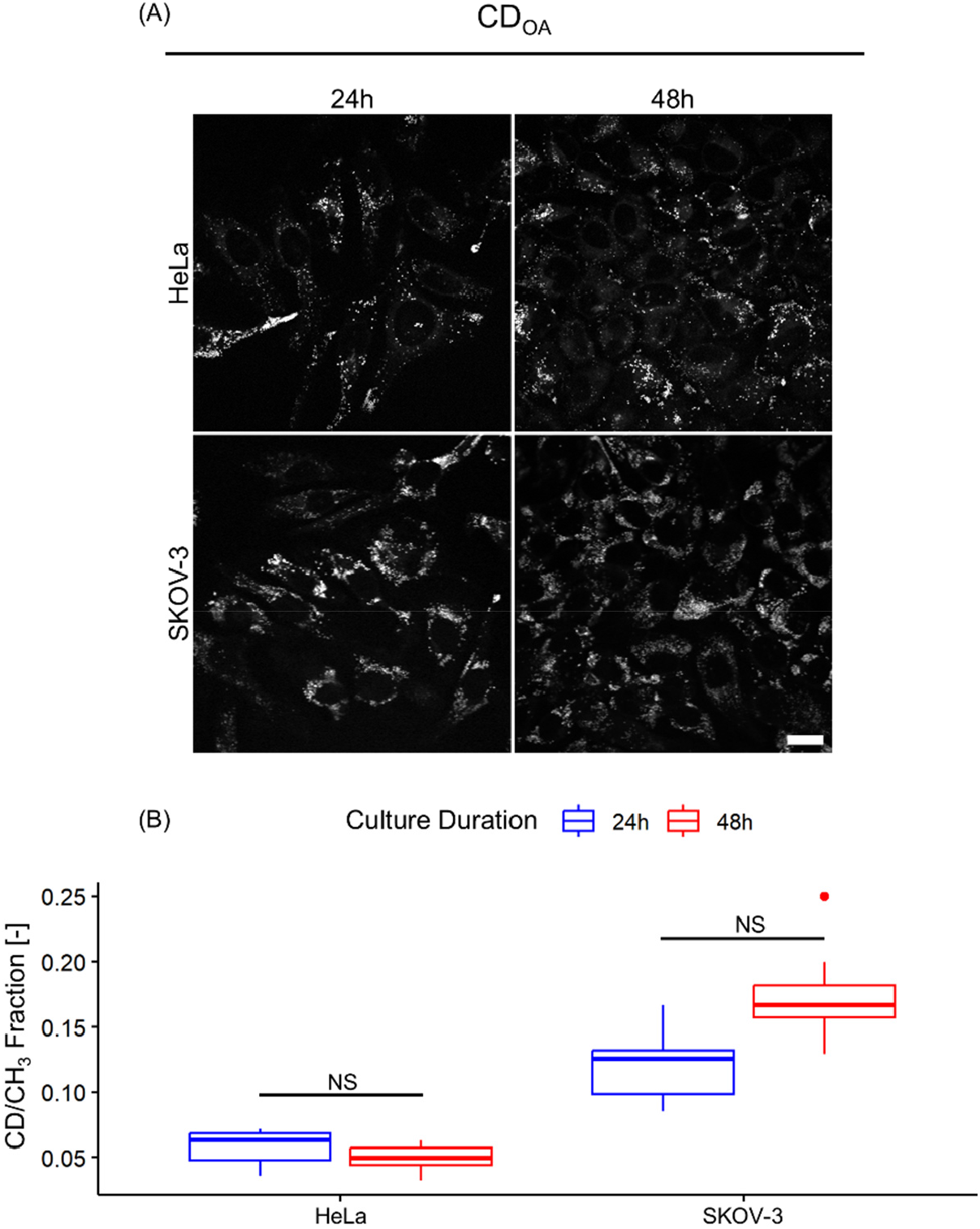
SKOV-3 and HeLa cells treated with 100 µM oleic acid-d_34_ for either 24 or 48 hours without medium renewal. n = 10 technical replicates for both 24h and 48h HeLa data. n = 9 technical replicates for 24h SKOV-3 data and n = 14 technical replicates for 48h SKOV-3 data.

**Fig. 6.**
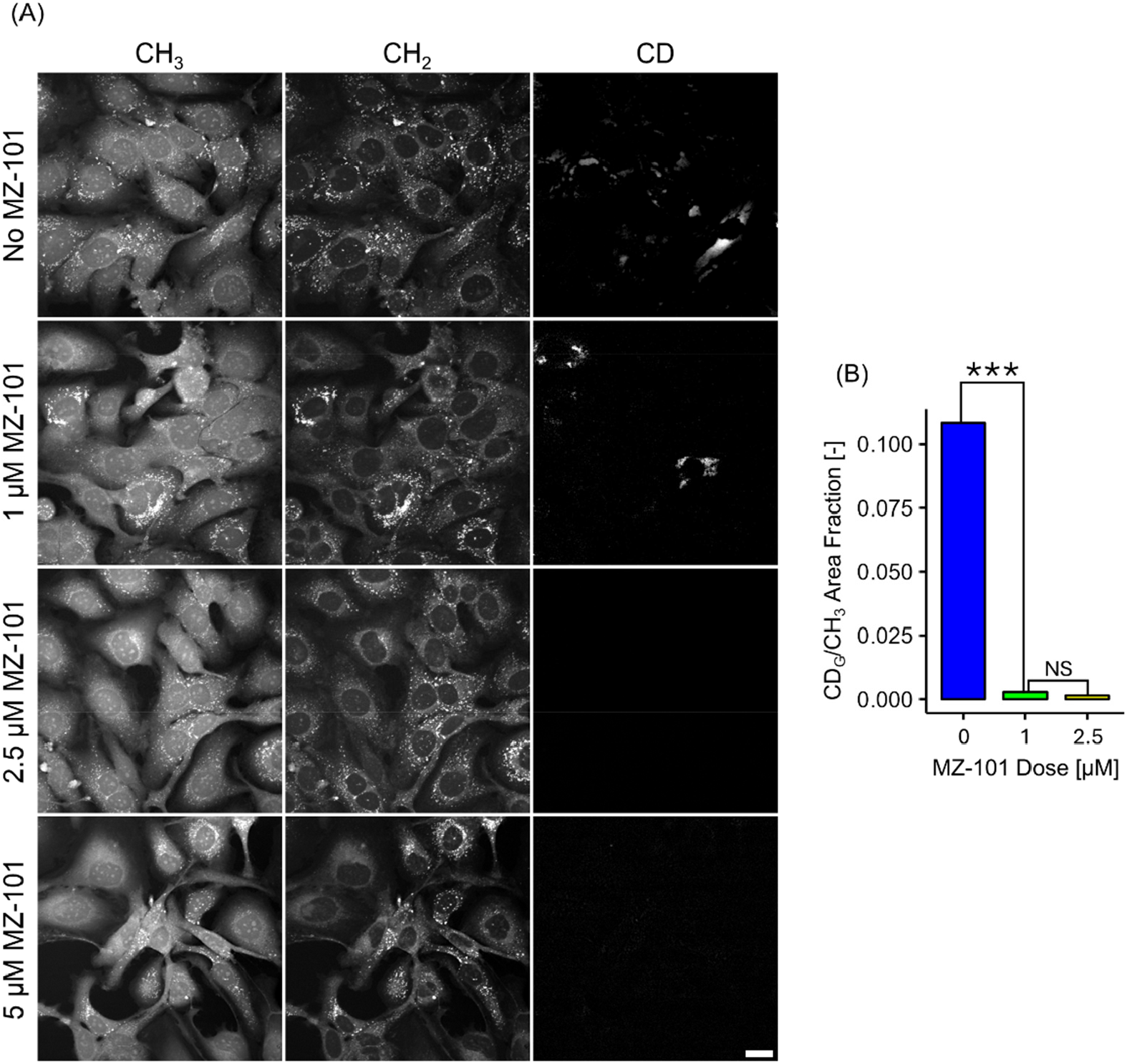
(A) Cell panel of live SKOV-3 cells treated with 25 mM D-glucose-d7 and GS-1 inhibitor (MZ-101) for over 72 hours. Scale bar = 20 µm. (B) CD area fraction quantification of MZ-101 impact on glycogen synthesis. Area fraction is reported as an average of 9 technical replicates. Statistical analyses were performed using one-way ANOVA followed by a Tukey’s HSD post hoc test. ‘^***^’ denotes a p value lower than 0.001. ‘NS’ denotes a non-significant result.

HeLa cells opting to reduce LD numbers over time may reflect a metabolic program geared toward rapid substrate mobilization in support of ongoing cell division. As nutrients are depleted from the medium, stored fatty acids can be liberated through lipolysis to maintain mitochondrial oxidation and biosynthetic pathways required for proliferation. This interpretation is consistent with reports that lipid droplet turnover provides an adaptive reservoir of energy-dense substrates for fast-dividing tumor cells [23].

Converse to the behavior observed in HeLa cells, the persistent accumulation of LDs in SKOV-3 cells implies a storage-focused metabolic phenotype. Rather than immediately mobilizing exogenous lipids for catabolism, SKOV-3 cells appear to prioritize the maintenance of lipid stores, which may act as a protective or buffering strategy against future metabolic stress. This aligns with prior work linking abundant lipid reserves to Ras-driven triple negative breast cancer cell survival under hypoxic or nutrient-poor conditions, as LDs not only provide an energy reserve but also sequester excess fatty acids to mitigate lipotoxicity [24].

When considered alongside the documented doubling times of the two cell lines, the observed differences in LD dynamics are biologically plausible. HeLa cells, with a shorter doubling time [25], have greater energetic and biosynthetic requirements per unit time, which may necessitate more aggressive mobilization of lipid stores. SKOV-3 cells, which divide more slowly, may instead benefit from sustaining stable lipid reserves over multiple cell cycles. Collectively, these findings suggest that SKOV-3 cells maintain a storage-centered metabolic profile, while HeLa cells rely more heavily on LD turnover to sustain rapid proliferation.

### 3.3 SKOV-3 subpopulations more efficiently polymerize glucosyl units into glycogen

To verify that the macromolecular species detected in SKOV-3 cells following treatment with deuterated glucose corresponded to glycogen, we employed a pharmacological approach targeting glycogen synthase-1 (GS-1), the key rate-limiting enzyme responsible for glycogen polymerization. A small-molecule inhibitor, MZ-101, was applied to cultured cells at graded concentrations to assess its impact on glycogen accumulation. At a concentration of 1 µM, a marked suppression of glycogen synthesis was observed, while treatment with 2.5 µM yielded near-complete abrogation of glycogen signal intensity (Fig. 8B). However, exposure to 5 µM MZ-101 resulted in discernible alterations in SKOV-3 cell morphology, characterized by an abundance of strand-like cells (Fig. 8A). This suggests a potential stress response at higher doses of MZ-101, although the exact mechanisms that govern these changes in cytoskeletal structure are unclear and require further validation. These results indicate that the observed CD chemical contrast and its accumulation is dependent on GS-1 activity, supporting glycogen as the molecular identity of the deuterium-labeled structure.

The observed sensitivity of SKOV-3 cells to GS-1 inhibition also highlights the reliance of these ovarian carcinoma cells on glucose-derived carbon flux into glycogen stores, even in the context of an environment where glycolytic pathways remain active. This dual channeling of glucose into both glycolytic and glycogenic pathways may reflect a strategy for maintaining metabolic flexibility under nutrient-rich conditions. Such a strategy would align with prior reports in other cancers that glycogen reserves can be mobilized to sustain proliferation during periods of environmental stress, including hypoxia or nutrient deprivation [26].

Moreover, analogous to the accumulation of lipid droplets, which has been proposed as a metabolic hallmark of aggressive cancer cell subsets, enhanced glycogen storage may serve as a parallel marker of tumor aggressiveness. By buffering fluctuations in nutrient supply, glycogen-rich cancer cells could maintain biosynthetic and energetic demands of cell cycle progression within the tumor microenvironment (TME). The heterogeneous expression pattern of glycogen observed in SKOV-3 cells is consistent with this model. Glycogen was not restricted to a bimodal expression pattern but instead distributed as a continuum of subcellular accumulation levels in untreated cultures. This heterogeneity became even more apparent following treatment with 1 µM MZ-101, where subpopulations of SKOV-3 cells exhibited reduced sensitivity to GS-1 inhibition, retaining measurable glycogen despite drug exposure. Such resistance underscores the potential for metabolic diversity within clonal cancer cell populations, which may complicate therapeutic targeting strategies. Together, these findings validate glycogen as the primary macromolecule detected in SKOV-3 cells following deuterated glucose labeling and highlight the potential biological significance of glycogen metabolism in supporting tumor cell adaptability and heterogeneity.

#### Lipid droplet depletion is more rapid in HeLa relative to SKOV-3

To further evaluate the capacity of HeLa cells to mobilize fatty acid–derived lipid substrates during nutrient stress, we quantified lipid droplet (LD) depletion dynamics in HeLa and SKOV-3 cells under progressive glucose starvation. Following a 24 h preloading period with 100 µM oleic acid (OA), cells were subjected to glucose depletion for 0–48 h, and LD abundance was assessed by BODIPY 493/503 fluorescence imaging. In HeLa cells, LDs were sparsely distributed under basal (0 h) glucose starvation and nearly absent after 48 h (Fig. 7A). Quantitative analysis confirmed a pronounced decrease in LD content relative to SKOV-3 cells, supporting the notion that HeLa cells exhibit enhanced lipid droplet hydrolysis, potentially reflecting greater reliance on neutral lipid catabolism to sustain proliferative metabolism (Fig. 7B). After 24 h of glucose starvation, HeLa cells consumed over 50% of accumulated LDs compared to non-starved controls, with a modest but nonsignificant additional decrease observed at 48 h. In contrast, cells only given growth medium instead of OA during the preloading phase displayed near-complete LD depletion after 24 h of glucose deprivation.

**Fig. 7.**
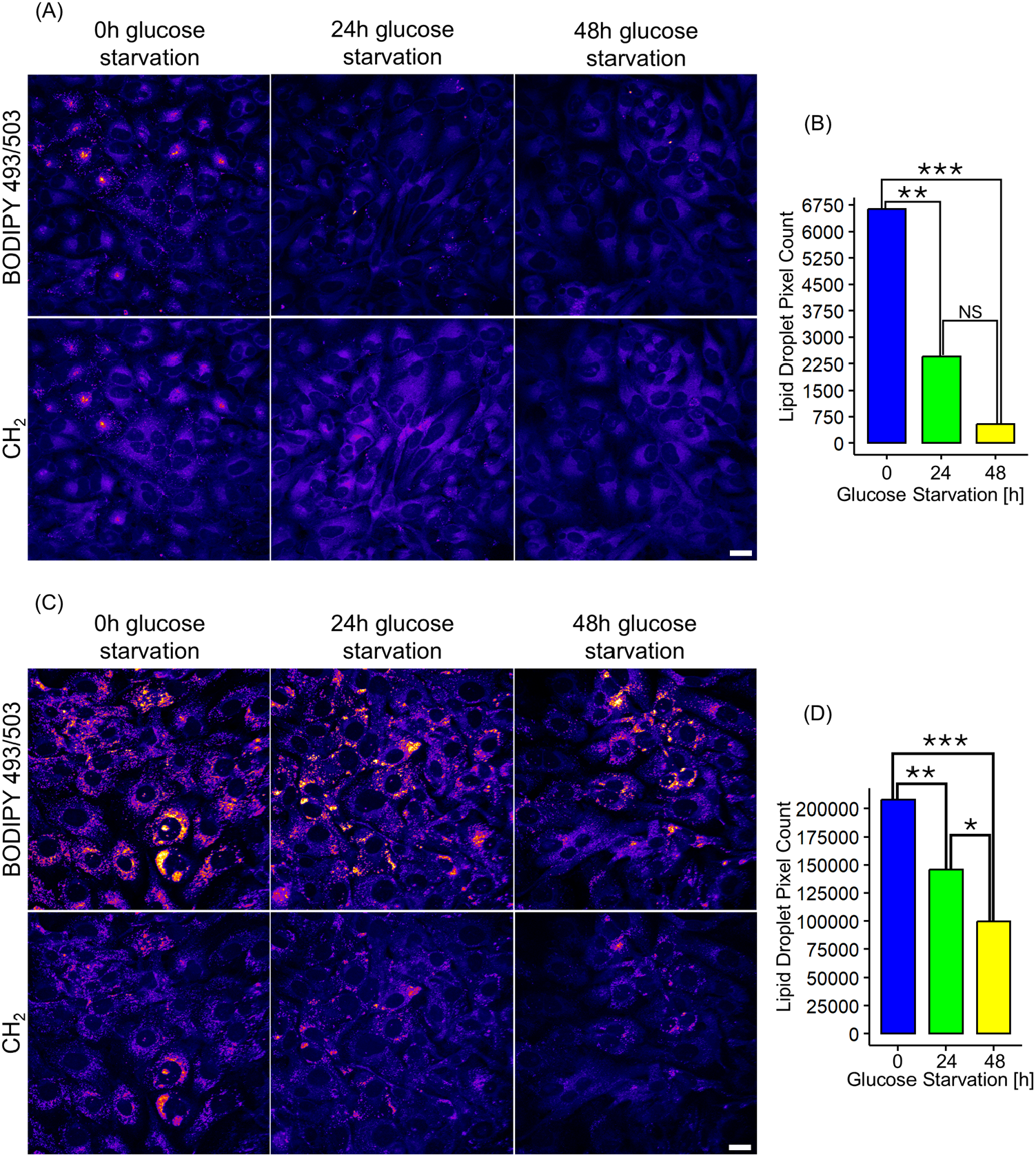
(A) SRS images at 2854 cm^-1^ and TPF images of BODIPY 493/503 stained OA loaded HeLa cells given no period of glucose starvation, 24h of glucose starvation, and 48h glucose starvation, respectively. (B) Quantification and statistical analysis of LD pixel area for each culture condition. (C) SRS images at 2854 cm^-1^ and TPF images of BODIPY 493/503 stained OA loaded SKOV-3 cells given no period of glucose starvation, 24h of glucose starvation, and 48h glucose starvation, respectively. (D) Quantification and statistical analysis of LD pixel area for each culture condition. Scale bar = 20 µm. ‘^***^’ denotes a p value lower than .001. ‘^**^’ denotes a p value lower than .01. ‘^*^’ denotes a p value lower than .05. ‘NS’ denotes a non-significant result. n = 5 technical replicates for each condition.

In contrast to HeLa cell behavior, LDs in all culture groups apart from the 48h glucose starvation condition for SKOV-3 showed a uniform distribution in most of the cell population, and in some locations, LDs were densely distributed intracellularly, contributing to elevated levels of both BODIPY 493/503 and CH_2_ signal (Fig. 7C). Following LD quantification, SKOV-3 cells demonstrated observable LD depletion, but at a markedly reduced rate. Compared to the non-glucose starved culture control, 24h of glucose starvation had only consumed approximately 33% of accumulated LDs. Interestingly, LD depletion rate in SKOV-3 cells as a function of glucose starvation time nearly followed an inverse linear relationship, although additional biological replicates would need to validate this behavior.

## 4. Discussion

Epithelial ovarian cancer continues to be one of the deadliest cancer diagnoses for women, with an estimated 5-year survival rate of 29% for late-stage diagnoses [27]. The demand for personalized care approaches for a patient is increasing, as it has become clear that every tumor has a unique and complex landscape of stromal cells, infiltrating immune cells, and a carefully regulated metabolome that can contribute to cancer progression and metastasis. It is therefore pertinent that efforts are put towards the discovery of new methods to characterize the TME and its constituents. The phenotyping of patient tumors using chemically informative techniques has potential to perform more rapid, non-invasive diagnostics. This pursuit is seen in the development of various Raman-based endoscopic probes, although more efforts are needed before translation to clinical use [28-30].

Here, we phenotype fatty acid and glycogen metabolism in epithelial ovarian cancer and cervical cancer cell line models. Glycogen dynamics in melanoma cells have been previously demonstrated using SRS with deuterium labeling [31], yet compared to other modes of cellular metabolism, its role in tumor progression remains understudied. The intracellular distribution and synthesis efficiency of produced deuterated macromolecules differed greatly between the two models, with SKOV-3 demonstrating greater single cell diversity regarding glycogen regulation compared to HeLa. Consequently, further work should emphasize the need to profile single cell metabolism in metabolically diverse cancer pathologies. This work also determined key differences in LD synthesis and turnover under nutrient starvation between HeLa and SKOV-3 cell lines, pointing towards LDs as a potentially diagnostic marker for cancer pathologies. SRS microscopy combined with deuterium labeling of metabolites can serve as a method to screen for metabolic diversity in cancer cell subpopulations, with utilization of the silent region of the Raman spectrum providing superior chemical contrast in biological samples. Owing to the reprogramming of cancer cells in response to changes in the TME metabolome, combining anti-cancer therapies with metabolic disruption could also be a promising strategy to improve patient survival.

## 5. Conclusion

This study demonstrates that SRS microscopy with deuterium-labeled metabolites can sensitively resolve metabolic heterogeneity in cancer cells. SKOV-3 and HeLa models exhibited distinct behaviors in both glycogen synthesis and lipid droplet turnover, with SKOV-3 showing greater single-cell variability in glycogen regulation. These findings highlight the importance of profiling metabolism at single-cell resolution to capture differences across cancer types. Our results further suggest that lipid storage dynamics may serve as useful metabolic markers for distinguishing cancer pathologies. Overall, silent-region SRS imaging provides a powerful platform for metabolic phenotyping and may help inform diagnostic strategies and therapeutic approaches that target metabolic vulnerabilities.

## References

1. O. Warburg, “On the origin of cancer cells,” Science 123, 309–314 (1956).

2. Z. Wang, et al., “Lipid metabolism as a target for cancer drug resistance: progress and prospects,” Front Pharmacol 14, 1274335 (2023).

3. A. Sebestyen, et al., “The role of metabolic ecosystem in cancer progression - metabolic plasticity and mTOR hyperactivity in tumor tissues,” Cancer Metastasis Rev 40, 989–1033 (2021).

4. Z. Xiao, et al., “Metabolic landscape of the tumor microenvironment at single cell resolution,” Nat Commun 10, 3763 (2019).

5. N. Ramamonjisoa and E. Ackerstaff, “Characterization of the Tumor Microenvironment and Tumor-Stroma Interaction by Non-invasive Preclinical Imaging,” Front Oncol 7, 3 (2017).

6. S. Chaudhry, et al., “Targeting lipid metabolism in the treatment of ovarian cancer,” Oncotarget 13, 768–783 (2022).

7. N. Koundouros and G. Poulogiannis, “Reprogramming of fatty acid metabolism in cancer,” Br J Cancer 122, 4–22 (2020).

8. Y. Matsushita, et al., “Lipid Metabolism in Oncology: Why It Matters, How to Research, and How to Treat,” Cancers (Basel) 13(2021).

9. K. Adekola, et al., “Glucose transporters in cancer metabolism,” Curr Opin Oncol 24, 650–654 (2012).

10. I. Barba, et al., “Targeting the Warburg Effect in Cancer: Where Do We Stand?,” Int J Mol Sci 25(2024).

11. D. Mishra and D. Banerjee, “Lactate Dehydrogenases as Metabolic Links between Tumor and Stroma in the Tumor Microenvironment,” Cancers (Basel) 11(2019).

12. Q. Gao and J. M. Goodman, “The lipid droplet-a well-connected organelle,” Front Cell Dev Biol 3, 49 (2015).

13. L. L. Listenberger, et al., “Triglyceride accumulation protects against fatty acid-induced lipotoxicity,” Proc Natl Acad Sci U S A 100, 3077–3082 (2003).

14. M. Pan, et al., “Lipid Metabolism and Lipidomics Applications in Cancer Research,” Adv Exp Med Biol 1316, 1–24 (2021).

15. L. M. Butler, et al., “Lipids and cancer: Emerging roles in pathogenesis, diagnosis and therapeutic intervention,” Adv Drug Deliv Rev 159, 245–293 (2020).

16. R. R. Jones, et al., “Raman Techniques: Fundamentals and Frontiers,” Nanoscale Res Lett 14, 231 (2019).

17. L. Wei, et al., “Live-Cell Bioorthogonal Chemical Imaging: Stimulated Raman Scattering Microscopy of Vibrational Probes,” Acc Chem Res 49, 1494–1502 (2016).

18. Y. Yuan, et al., “Visualizing drug-induced lipid accumulation in lysosomes of live cancer cells with stimulated Raman imaging,” Biomed Opt Express 14, 2551–2564 (2023).

19. M. Tenopoulou, et al., “Does the calcein-AM method assay the total cellular ‘labile iron pool’ or only a fraction of it?,” Biochem J 403, 261–266 (2007).

20. M. Kopec, et al., “Hyperglycemia and cancer in human lung carcinoma by means of Raman spectroscopy and imaging,” Sci Rep 12, 18561 (2022).

21. J. A. Menendez and R. Lupu, “Fatty acid synthase and the lipogenic phenotype in cancer pathogenesis,” Nat Rev Cancer 7, 763–777 (2007).

22. K. M. Nieman, et al., “Adipocytes promote ovarian cancer metastasis and provide energy for rapid tumor growth,” Nat Med 17, 1498–1503 (2011).

23. S. Rambold, et al., “Fatty acid trafficking in starved cells: regulation by lipid droplet lipolysis, autophagy, and mitochondrial fusion dynamics,” Dev Cell 32, 678–692 (2015).

24. E. Jarc, et al., “Lipid droplets induced by secreted phospholipase A(2) and unsaturated fatty acids protect breast cancer cells from nutrient and lipotoxic stress,” Biochim Biophys Acta Mol Cell Biol Lipids 1863, 247–265 (2018).

25. S. P. Langdon, “Characterization and authentication of cancer cell lines: an overview,” Methods Mol Med 88, 33–42 (2004).

26. E. Favaro, et al., “Glucose utilization via glycogen phosphorylase sustains proliferation and prevents premature senescence in cancer cells,” Cell Metab 16, 751–764 (2012).

27. L. C. Peres, et al., “Invasive Epithelial Ovarian Cancer Survival by Histotype and Disease Stage,” J Natl Cancer Inst 111, 60–68 (2019).

28. L. Zavaleta, et al., “A Raman-based endoscopic strategy for multiplexed molecular imaging,” Proc Natl Acad Sci U S A 110, E2288–2297 (2013).

29. A. Lombardini, et al., “High-resolution multimodal flexible coherent Raman endoscope,” Light Sci Appl 7, 10 (2018).

30. P. Zirak, et al., “Invited Article: A rigid coherent anti-Stokes Raman scattering endoscope with high resolution and a large field of view,” APL Photonics 3(2018).

31. D. Lee, et al., “Visualizing Subcellular Enrichment of Glycogen in Live Cancer Cells by Stimulated Raman Scattering,” Anal Chem 92, 13182–13191 (2020).

